# West Nile virus detection in horses in three Brazilian states

**DOI:** 10.1101/2021.01.06.425363

**Authors:** Erica Azevedo Costa, Marta Giovanetti, Lilian Silva Catenacci, Vagner Fonseca, Flavia Aburjaile, Felipe Melo Campos Iani, Marcelo Adriano da Cunha e Silva Vieira, Danielle Freitas Henriques, Daniele Barbosa de Almeida, Maria Isabel Maldonado Coelho Guedes, Beatriz Senra Álvares da Silva Ramos, Aila Solimar Gonçalves Silva, Tulio de Oliveira, Karina Ribeiro Leite Jardim Cavalcante, Noely Fabiana Oliveira de Moura, Alessandro Pecego Martins Romano, Carlos F. Campelo de Albuquerque, Lauro César Soares Feitosa, José Joffre Martins Bayeux, Raffaella Bertoni Cavalcanti Teixeira, Osmaikon Lisboa Lobato, Silvokleio da Costa Silva, José Lourenço, Luiz Carlos Junior Alcantara

**Affiliations:** Departamento de Medicina Veterinária Preventiva, Escola de Veterinaria, Universidade Federal de Minas Gerais, Belo Horizonte, Minas Gerais, Brazil; Laboratório de Flavivírus, Instituto Oswaldo Cruz, Fundação Oswaldo Cruz, Rio de Janeiro, Brazil; Laboratório de Genética Celular e Molecular, Universidade Federal de Minas Gerais, Belo Horizonte, Minas Gerais, Brazil; Departamento De Morfofisiologia Veterinária, Universidade Federal do Piauí; KwaZulu-Natal Research Innovation and Sequencing Platform (KRISP), School of Laboratory Medicine and Medical Sciences, College of Health Sciences, University of KwaZulu-Natal, Durban, South Africa; Coordenação Geral dos Laboratórios de Saúde Pública/Secretaria de Vigilância em Saúde, Ministério da Saúde, (CGLAB/SVS-MS) Brasília, Distrito Federal 70719-040, Brazil; Laboratório Central de Saúde Pública, Fundação Ezequiel Dias, Belo Horizonte, Brazil; Piaui State Health Department, Terezina, Brazil; Seção de Arbovirologia e Febres Hemorrágicas, Instituto Evandro Chagas, Ministério da Saúde; Coordenacao Geral das Arboviroses, Secretaria de Vigilância em Saúde/Ministério da Saúde, Brasília, Distrito Federal, Brazil; Organização PanAmericana da Saúde/Organização Mundial da Saúde, Brasília-DF, Brazil; Departamento de Clínica e Cirurgia Veterinária, Centro de Ciências Agrárias, Universidade Federal do Piauí, Teresina, Piauí, Brazil; UNIVAP-Universidade Vale do Paraíba, Faculdade de Ciências da Saúde, Medicina Veterinária, Urbanova, São José Dos Campos, SP; Departamento de Clinica e Cirurgia Veterinárias, Escola de Veterinaria, Universidade Federal de Minas Gerais, Belo Horizonte, Minas Gerais, Brazil; Laboratório de Genética e Conservação de Germoplasma, Campus Prof.^a^ Cinobelina Elvas, Universidade Federal do Piauí, Bom Jesus, Piauí, Brazil; Department of Zoology, University of Oxford, Oxford OX1 3PS, UK

**Keywords:** West Nile virus, genomic monitoring, molecular detection, Brazil

## Abstract

We report genetic evidence of WNV circulation from southern and northeastern Brazilian states isolated from equine red blood cells. In the northeastern state the tenth human case was also detected, presenting neuroinvasive disease compatible with WNV infection. Our analyses demonstrate that much is still unknown on the virus’ local epidemiology. We advocate for a shift to active surveillance, to ensure adequate control for future epidemics with spill-over potential to humans.

## Text

West Nile virus (WNV), a member of the *Flaviviridae* family, was first identified in the West Nile district of Uganda in 1937, but is nowadays commonly found in Africa, Europe, North America, the Middle East, and Asia [1–3].

WNV transmission is maintained in a mosquito-bird cycle, for which the genus *Culex,* in particular *Cx. pipiens,* are considered the principal vectors [4]. WNV can infect humans, equines and other mammals, but these are considered “dead-end” hosts, given their weak potential to function as amplifying hosts to spread infection onwards [5, 6]. Around 80% of WNV infections in humans are asymptomatic while the rest may develop mild or severe disease. Mild disease includes fever, headache, tiredness and vomiting [7, 8], while severe disease (neuroinvasive) is characterized by high fever, coma, convulsions and paralysis [7, 8]. Equine infections can occasionally cause neurological disease and death [7, 8], such that equines typically serve as sentinel species for WNV outbreaks with potential for spill-over into human populations.

Genome detection of WNV in South America were originally reported in horses (Argentina in 2006) and captive flamingos (Colombia, in 2012) [9, 10]. The first ever sequenced genome in Brazil was in 2019, when the virus was isolated from a horse sicked with severe neurological disease in the Espírito Santo state [11]. Despite multiple studies reporting serological evidence suggestive of WNV circulation in Brazil [11–13] and reports human WNV disease confirmed cases in the Piauí state [13], much is unknown about genomic diversity, evolution and transmission dynamics across the country.

Here, we report genetic evidence of WNV circulation in three Brazilian states extracted from equine red blood cells (RBCs) with neurological or ophthalmic disease and use a computational approach to explore the theoretical transmission potential of WNV within one of those states.

Samples (RBCs) from three horses with suspected WNV infection obtained from southern (Minas Gerais and São Paulo) and northeastern (Piauí) Brazilian states were sent for molecular diagnosis at the *Departamento de Medicina Veterinária Preventiva* at the Federal University of Minas Gerais (UFMG) (**for details see Appendix**).

RNAs were extracted from red blood cells and tested using an in-house PCR assay (**see Technical Appendix for details**). WNV-specific RT-PCR amplification products were obtained using a combination of nested and multiplex PCR scheme (**Figure 1 panel A, B**) (**see Technical Appendix for details**). A multiplex PCR primer scheme was then designed (**Appendix Table S1**) to generate complete genomes sequences by means of portable nanopore sequencing. The published WNV genome from Brazil (MH643887) was used to generate a mean 98.4% consensus sequences that formed the target for primer design. New genomes were deposited in the GenBank with accession numbers MW420987, MW420988 and MW420989 (**Table 1).**

**Figure 1.**
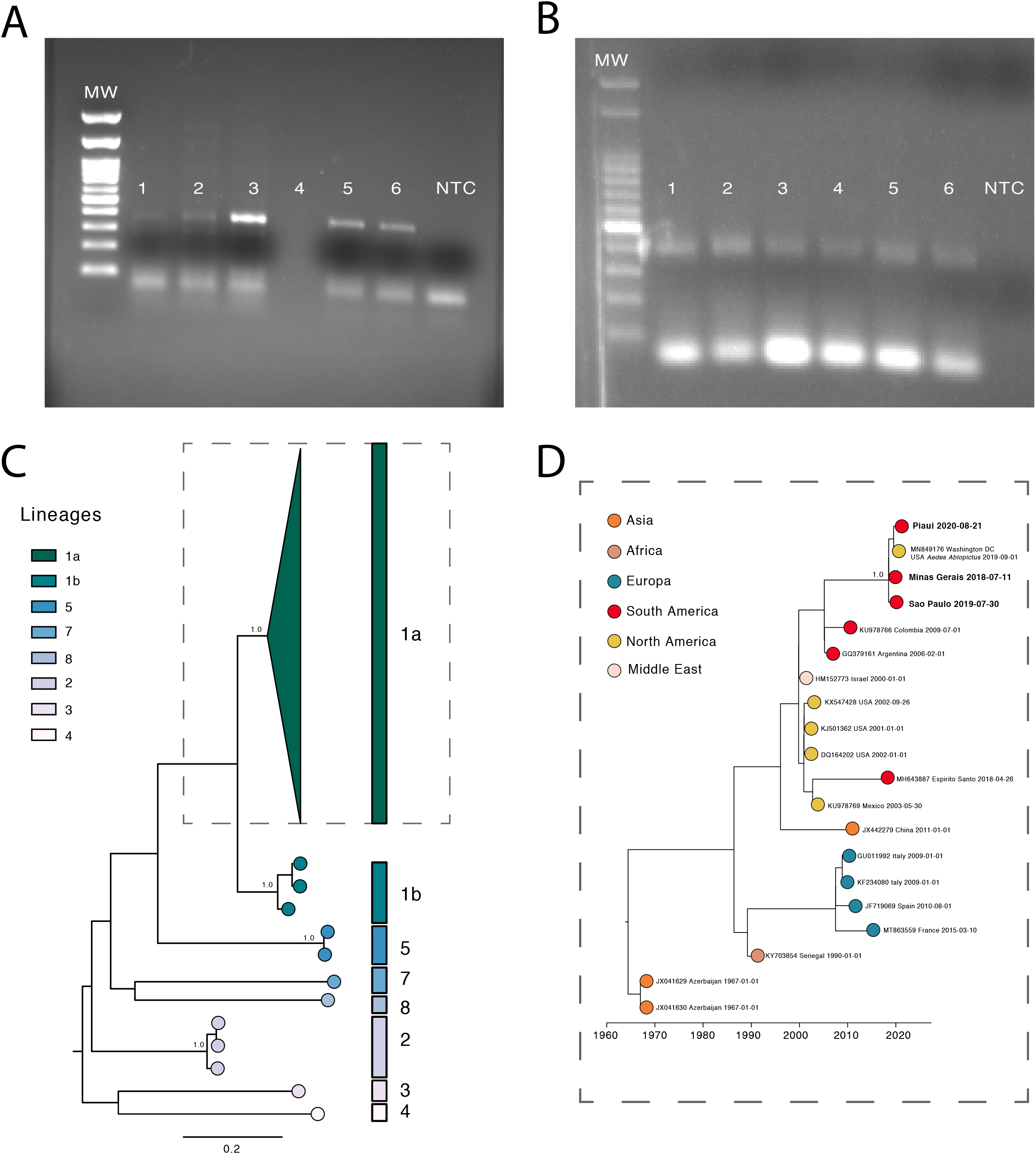
Investigation of WNV infections in Brazil, between July 2018 – September 2020. A-B) Agarose gel electrophoresis of amplicons from assay for WNV. A) nested RT-PCR. MW-Molecular weight marker; 1-plasm of horse from São Paulo; 2-buffy coat of horse from São Paulo; 3-washed RBC of horse from São Paulo; 4-without sample; 5 and 6-positive control (synthetic gene); NTC, no template control. (B) Multiplex PCR. MW-Molecular weight marker; 1-horse form Minas Gerais (pair primers); 2-horse form Minas Gerais (impair primers); 3-horse form Sao Paulo (pair primers); 4-horse form Sao Paulo (impair primers); 5-horse form Piaui (pair primers); 6-horse form Piaui (impair primers); NTC, no template control. C) Midpoint rooted maximum-likelihood phylogeny of WNV genomes, showing major lineages. The scale bar is in units of substitutions per site (s/s). Support for branching structure is shown by bootstrap values at nodes. D) Time-resolved maximum likelihood tree showing the WNV strains belonged to the 1a lineage. Colors indicate geographic location of sampling. The new Brazilian WNV strains are shown in bold.

**Table 1.** Epidemiological information and sequencing statistics of the 3 sequenced samples of WNV sampled in Minas Gerais, Sao Paulo, Piaui Brazilian states.

| ID | Sample | Collection date | Age | Sex | State | Municipality | Reads | Coverage (%) | Depth of coverage | Lineage Assignment | Accession Number | Clinical sign |
| --- | --- | --- | --- | --- | --- | --- | --- | --- | --- | --- | --- | --- |
| BC02_07 | RBCs | 11/07/2018 | 9 months | F | MG | Sabara | 343743 | 97.9 | 6527.6 | Lineage 1a | MW420989 | Chorioretinitis |
| BC03_04 | RBCs | 30/07/2019 | 13 years-old | M | SP | São Bernardo do Campo | 170980 | 97.9 | 3189.7 | Lineage 1a | MW420988 | Muscle stiffness, tremor retinal and flaccid paralysis |
| BC05_06 | RBCs | 21/08/2020 | 5 years-old | F | PI | Parnaíba | 222516 | 99.4 | 4121.4 | Lineage 1a | MW420987 | Neurological complications |

ID=study identifier; RBCs=Red Blood Cells; Collection date=Sample collection date; Municipality=Municipality of residence; State= MG-Minas Gerais; SP-Sao Paulo; PI-Piaui; Sex: M=Male; F=Female; Accession Number=NCBI accession number

We constructed phylogenetic trees to explore the relationship of the sequenced genomes to those from elsewhere globally. We retrieved 2321 WNV genome sequences with associated lineage date and country of collection from GenBank, from which we generated a subset that included the highly supported (>0.9) clade containing the newly WNV strains obtained in this study plus 29 sequences (randomly sampled) from all lineages and performed phylogenetic analysis (**see Technical Appendix for detail**). An automated online phylogenetic tool to identify and classify WNV sequences was developed (available at http://krisp.ukzn.ac.za/app/typingtool/wnv/job/9b40f631-51c4-419c-9edf-2206e7cd8d9c/interactive-tree/phylo-WNV.xml).

Phylogenies estimated by the newly developed WNV typing tool, along with maximum likelihood methods (**Figure 1 panel C),** consistently placed the Brazilian genomes in a single clade within the 1a lineage with maximum statistical support (bootstrap = 100%) (**Supplementary Figure 1**).

Time-resolved maximum likelihood tree appeared to be consistent with previous estimates [11] and showed that the new genomes clustered with strong bootstrap support (97%) with a WNV strain isolated from an *Aedes albopictus* mosquito in Washington DC, USA in 2019 (**Figure 1 panel D**). Interestingly, these new isolates did not group with the previously sequenced genome in 2019 from the Espirito Santo state, suggesting that inter-continental introduction events might be frequent in Brazil.

We also explored the dynamics of suspected human WNV cases (**see Technical Appendix for detail**) in the three Brazilian states for which we had sequence data. Between late 2015 and early 2020, all states had suspected cases, suggesting continuous circulation or recent importation from other regions of the country or elsewhere. The state of Piauí presented sufficient reports (N=116) for an exploration of the geo-temporal dynamics of WNV spread (**Figure 2A**), while the other two states had a much smaller number of total reports (N=3 for MG, 18 for SP).

**Figure 2.**
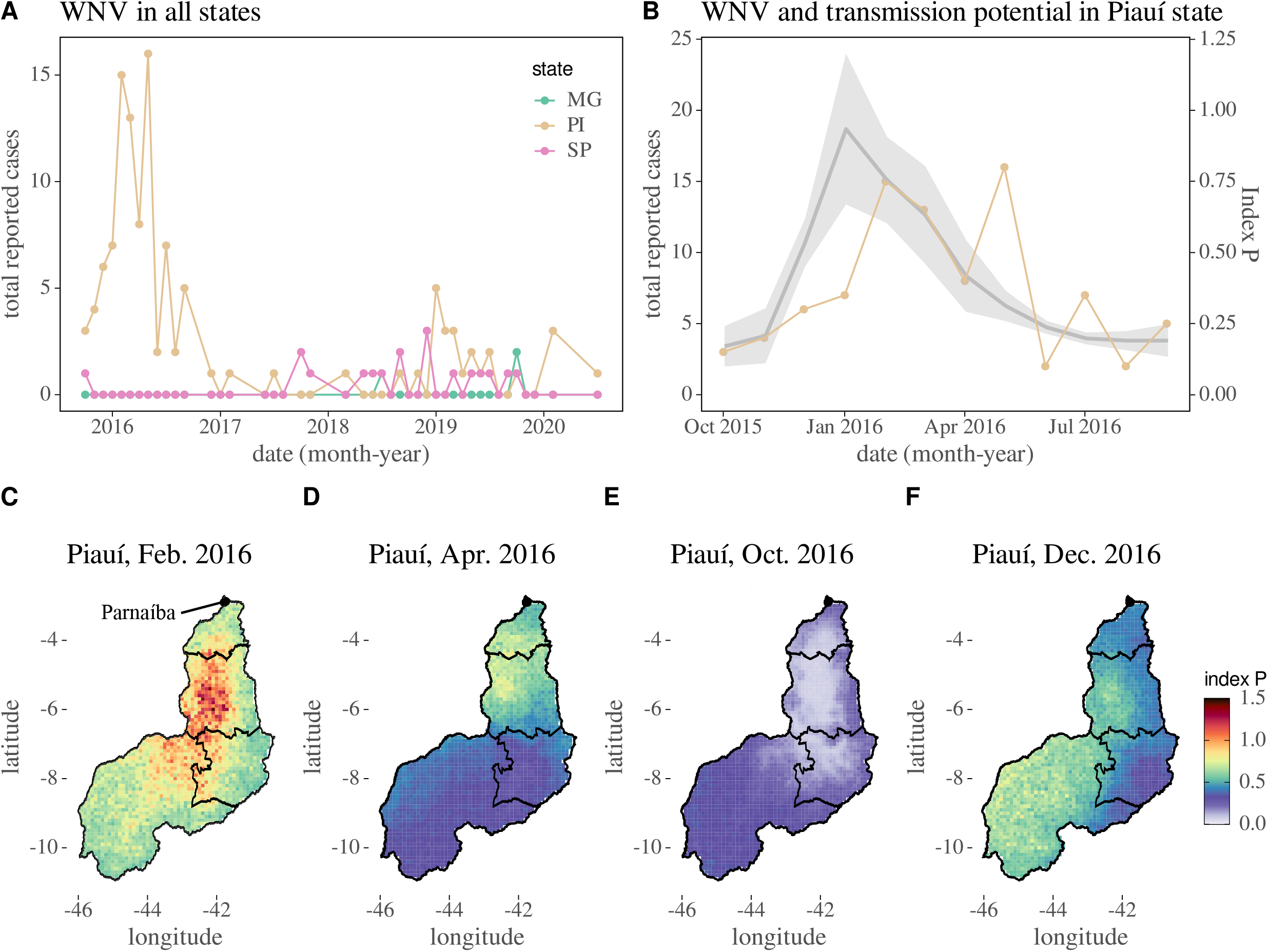
WNV reporting of suspected cases and estimated transmission potential. (A) Time series of suspected human WNV cases in the Brazilian states of Minas Gerais (MG, green), São Paulo (SP, pink) and Piauí (PI, gold). (B) WNV human suspected cases and estimated monthly mean transmission potential (index P, grey) for the state of Piauí (gold). The shaded area includes the standard deviation. (C-F) Maps of estimated transmission potential (index P) per geo-location for four different months in the year of 2016 (from C to F, February, April, October, December respectively). Color legend on the right applies to all maps. The city of Parnaíba from where the Piauí sequence is available is marked by a black, full point in the north of the state.

To estimate the transmission potential of WNV we calculated the index P, a computational approach informed by climatic variables from Lourenço et al. recently applied in Israel [14] (**see Technical Appendix for detail**). In the 2015-2016 season, cases in Piauí presented a seasonal signal typical of WNV and other mosquito-borne viruses, with a peak between February and May (late summer and autumn). We compared suspected cases per month to the monthly mean transmission potential across the state (**Figure 2B**). The Spearman’s correlation between the two variables was 0.66 (p-value 0.01). This correlation was smaller than previously reported for Israel (>0.9), likely resulting from the suspected nature of the Brazillian cases, or from averaging climatic variables (and thus the index P) across the vast spatial dimension of the state.

We also estimated the index P in space for this season, which highlighted some geo-patterns (**Figures 2C-F**). In summer, when reports were high, estimated transmission potential was mainly highest in the center at the −43 longitude axis (**Figure 2C).** Into autumn, the southern region was the first with lower transmission potential (**Figure 2D**), followed by the northern region in the winter (**Figure 2E**). In spring, coinciding with the trough of reporting, the lowest transmission potential was estimated in the same central region presenting highest potential in the summer and autumn (**Figure 2F**). These geo-temporal snapshots highlight a possible wave of seasonal transmission potential, starting in the south-west in spring (**Figure 2F**), moving to the north-east in summer (**Figure 2C**), and ending in the north during autumn (**Figure 2D**).

## Conclusions

Our analyses indicate that additional data is required to better identify routes of WNV importation into Brazil, and to understand its local transmission dynamics. Critically, it is still uncertain where in the country WNV is endemic. Indeed, if under reporting is frequent, the data presented in this article is compatible with both sporadic or endemic local transmission. Furthermore, the detection of WNV RNA from equine whole blood presented in this study proved to be an effective diagnostic method in horses. Shifting from passive to active WNV screening and sequencing in equines and birds in Brazil must be implemented to better understand the virus’ local epidemiology, and to be able to act accordingly to prevent any future epidemics with significant spill-over to humans.

## Acknowledgements

The authors thank the important contributions of the Municipal and Piaui State Health Department (SESAPI, FMS), Municipal and Piaui State Animal Health Department (ADAPI), Laboratório de Saúde Pública do Piauí (LACEN-PI) and the colleague Thiago dos Santos Silva. We also thank the sponsoring institutions: Saint Louis Zoo WildCare Institute and Institute for Conservation Medicine (USA), Universidade Federal do Piauí (UFPI), Fundação de Amparo a Pesquisa do Estado do Piauí (FAPEPI). Authors also thank the Municipal and State Health Department of São Paulo and Minas Gerais state.

## Data Sharing

Newly generated WNV sequences have been deposited in GenBank under accession numbers MW420987, MW420988 and MW420989.

## References

[1] Fall G, Di Paola N, Faye M, et al. Biological and phylogenetic characteristics of West African lineages of West Nile virus (DWC Beasley, Ed.). PLoS Negl Trop Dis. 2017;11: 1–23.

[2] Smithburn KC, Hughes TP, Burke AW, et al. A neutrotropic virus isolated from the blood of a native of Uganda. Am J Trop Med Hyg. 1940; 20: 471–2.

[3] Murgue B, Zeller H, Deubel V. The ecology and epidemiology of West Nile virus in Africa, Europe and Asia. Curr Top Microbiol Immunol. 2002;267:195–221.

[4] Campbell GL, Marfin AA, Lanciotti RS, et al. West Nile virus. Lancet Infect Dis. 2002;2:519–29.

[5] Gamino V, Ho□fle U. Pathology and tissue tropism of natural West Nile virus infection in birds: a review. Vet Res. 2013;44:46–89.

[6] Bunning ML, Bowen RA, Cropp CB, et al. Experimental infection of horses with West Nile virus. Emerg Infect Dis. 2002;8:380–6.

[7] Hayes EB, Sejvar JJ, Zaki SR, et al. Virology, pathology, and clinical manifestations of West Nile virus disease. Emerg Infect Dis. 2005;11:1174–1179.

[8] Kramer LD, Li J, Shi P-Y. West Nile virus. Lancet Neurol. 2007;6:171–181.

[9] Morales MA, Barrandeguy M, Fabbri C, et al. West Nile virus isolation from equines in Argentina. Emerg Infect Dis. 2006;12:1559–61.

[10] Osorio JE, Ciuoderis KA, Lopera JG, et al. Characterization of West Nile viruses isolated from captive American flamingoes (Phoenicopterus ruber) in Medellin, Colombia. Am J Trop Med Hyg. 2012;87:565–72.

[11] Martins LC, Silva EVPD, Casseb LMN, et al. First isolation of West Nile virus in Brazil. Mem Inst Oswaldo Cruz. 2019 Jan 17;114:e180332.

[12] Pauvolid-Corre a A, Morales MA, Levis S, Figueiredo LTM, Cou-to-Lima D, Campos Z, et al. Neutralising antibodies for West Nile virus in horses from Brazilian Pantanal. Mem Inst Oswaldo Cruz. 2011;106:467–74.

[13] Vieira MA, Romano APM, Borba AS, et al. West Nile Virus Encephalitis: The First Human Case Recorded in Brazil. Am J Trop Med Hyg. 2015; 93(2):377–9.

[14] Lourenço J, Thompson RN, Thézé J, et al. Characterising West Nile virus epidemiology in Israel using a transmission suitability index. Euro Surveill. 2020;2:5–41.

