## Appendix for "West Nile virus detection in horses in three Brazilian states"

**Technical Appendix**

**Materials and Methods**

**Ethics statement**

This project was reviewed and approved by the Comissão Nacional de Ética em Pesquisa (CONEP) [National Research Ethics Committee] from the Brazilian Ministry of Health (BrMoH), as part of the arboviral genomic surveillance efforts within the terms of Resolution 510/2016 of CONEP, by the Pan American Health Organization Ethics Review Committee (PAHOERC) (Ref. No. PAHO-2016-08-0029), by the Animal Welfare Committee of Universidade Federal do Piauí, under n°065/19 and by the Oswaldo Cruz Foundation Ethics Committee (CAAE: 90249218.6.1001.5248).

**Sample collection, viral RNA isolation and PCR screening**

Samples (red blood cells - RBCs) from three horses with suspected WNV infection obtained from southern (Minas Gerais and São Paulo) and northeastern (Piauí) Brazilian states were sent for molecular diagnosis at the Laboratório de Patologia Molecular at the Federal University of Minas Gerais (UFMG).

Sample 1 from July 11^st^ 2018 was collected from a 9 months-old female horse in a farm in the state of Minas Gerais, Mangueiras neighbourhood (Sabará), 15 kilometres from the capital Belo Horizonte. Clinical findings were consistent with bilateral blindness. Neurological examination revealed no other abnormalities. The ophthalmological exams (direct and indirect pupillary light reflex-PLR, fluorescein eye stain test, fundus examination, intraocular pressure) were consistent with retinal disease, mainly with chorioretinitis.

Sample 2 from July 30^th^, 2019 was collected from a 13 years-old male horse who presented seizure episodes, muscle stiffness, tremor retinal and flaccid paralysis, in a farm located in São Bernardo do Campo countryside of the São Paulo state. Twenty-four days after the onset of neurological signs, the animal had severe pain in the forelimbs from laminitis and it was euthanized due to hoof decumulation.

Sample 3 from August 21^st^, 2020 was collected from a male horse, 5 years old, which died 72 hours after presenting neurological signs, in a farm located in the municipality of Parnaíba, Piauí state. The animal presented motor incoordination, paddling movements, loss of sensitivity over the spine column, and behavioral changes. In this municipality the tenth human case was also detected, presenting neuroinvasive disease compatible with WNV infection, confirmed by serological assay (IgM) in both serum and cerebrospinal fluid (CSF) samples during acute and convalescent phases.

Whole blood samples obtained from the three horses were centrifuged at 1260 x g for 20 min, and the plasma and buffy coat fractions were collected and stored at 4°C. Red blood cells (RBC) were washed by centrifugation three times in phosphate-buffered saline (PBS) at 1260 g for 10 min and stored also at 4°C [1]. RNA from each unit (washed RBC, plasma and buffy coat) were extracted using the QIAmp Viral RNA Mini kit (Qiagen), following manufacturer’s recommendations.

Diagnostic investigation of arboviruses was performed by a generic RT‐PCR targeting the flavivirus non-structural protein 5 (NS5) gene [2] and alphavirus non-structural protein 1 gene (nsP1) [3]. West Nile virus‐specific degenerated primers: forward primers (+) AACCKCCAGAAGGAGTSAAR and reverse primers (-) AGCYTCRAACTCCAGRAAGC were used in second reaction of nested PCR targeting the NS5 gene after a genus specific flavivirus RT-PCR amplification [4]. A synthetic gene fragment of partial NS5 gene (gblocks gene fragment, Integrated DNA Technologies) was used as a positive control. The 25 μl PCR “master‐mix” comprised 2.5 μl of 10× PCR buffer, 1.5 mM MgCl2, 0,4 μM of each primer (forward and reverse), 0.8 μM dNTP mixture (Phoneutria), 1 U Taq DNA polymerase (Platinum Taq DNA polymerase; Invitrogen), 2 μl of template DNA (sample or gBlock), and DNA/RNAse‐free water. The thermocycling conditions involved 40 cycles, and reaction conditions was previously reported in [4]. As an internal control for amplification efficiency, primers for the beta actin gene were used. As a negative control for the reactions, we used RNA extracted from equine washed RBC, plasma and buffy coat that previously tested negative for arboviruses, equine herpesvirus 1 and 4 and borna disease. The amplicons were analysed by 1% (w/v) agarose gel electrophoresis, stained with ethidium bromide and visualized under UV light. Nested PCR were performed for equine herpesvirus 1 (EHV-1) [5] and nested RT-PCR for borna disease [6], both with negative results in the 3 horses.

**cDNA synthesis and multiplex tiling PCR**

WNV positive (in nested RT-PCR) RNA samples from washed RBCs were then submitted to a cDNA synthesis protocol [*7*] using Superscript IV cDNA Synthesis Kit. Then, a multiplex tiling PCR was conducted using Q5 High Fidelity Hot-Start DNA Polymerase (New England Biolabs) and a WNV sequencing primers scheme (divided into two separated pools) designed using Primal Scheme (**Table S1**) ([http://primal.zibraproject.org](http://primal.zibraproject.org/)) [8]. The thermocycling conditions involved 40 cycles, and reaction conditions was previously reported in [8].

**Library preparation and nanopore sequencing**

Amplicons were purified using 1x AMPure XP Beads and cleaned-up PCR products concentrations were measured using Qubit™ dsDNA HS Assay Kit on a Qubit 3.0 fluorimeter (ThermoFisher). DNA library preparation was carried out using the Ligation Sequencing Kit and the Native Barcoding Kit (NBD104, Oxford Nanopore Technologies, Oxford, UK) [8]. Purified PCR products pools were pooled together before barcoding reactions (taking in consideration each amplicon pool DNA concentrations), and one barcode was used per sample in order to maximize the number of samples per flow cell. Sequencing library was loaded onto a R9.4 flow cell, and data was collected for up to 6 hours, but generally less.

**Generation of consensus sequences**

Raw files were basecalled using Guppy and barcode demultiplexing was performed using qcat. Consensus sequences were generated by *de novo* assembling using Genome Detective (<https://www.genomedetective.com/>app/) [9].

**West Nile Virus typing tool: Classification method and implementation**

The classification pipeline we present comprises two components. One for species and sub-species assignment that enables assignment at these levels by BLASTing the query sequences against a set reference sequences [10]. An assignment is made when BLAST reports a result that exceeds the present threshold.

The other component constructs a Neighbour Joining (NJ) phylogenetic tree that are used to make assignments at the lineages and sublineages level. For this component, the query sequence is aligned against a set of reference sequences using the profile alignment option in the ClustalW software [11], such that the query sequence is added to the existing alignment of reference sequences. Following the alignment, a NJ phylogenetic tree, with 100 bootstrap replicates is inferred. The tree is constructed using the HKY distance metric with gamma among-site rate variation, as implemented in the PAUP* software [12]. The query sequence is assigned to a particular genotype if it clusters monophyletically with that genotype clade with bootstrap support >70%. If the bootstrap support is <70%, the genotype is reported to be unassigned (**Supplementary Figure 1**).

For each of these steps, the earlier discussed reference strains were used, with respect to the appropriate typing level (i.e. virus species, lineages and sublineages). Testing revealed that a BLAST cut-off value of 200 allowed accurate identification of the virus species and WNV using sequence segments > 200 base pairs. Note that the species classification procedure is implemented as separate BLAST steps. This enables the tool to efficiently perform large throughput species classification, such as for the classification of shorts sequencing reads. An instance of the web application is publically available on a dedicated server (https://www.genomedetective.com/app/typingtool/wnv/). The web interface on this server accepts up to 2,000 whole-genome or partial genome sequences at a time.

**Phylogenetic analysis**

The 3 newly sequences reported in this study were initially submitted to a genotyping analysis using the new phylogenetic West Nile virus subtyping tool, available at <https://www.genomedetective.com/app/typingtool/wnv>. To put the newly WNV sequences in a global context, we constructed phylogenetic trees to explore the relationship of the sequenced genomes to those of other isolates.

We retrieved 2321 WNV genome sequences with associated lineage date and country of collection from GenBank (**Supplementary Figure 2**). From this dataset, we generated a subset that included the highly supported (>0.9) clade containing the newly WNV strains obtained in this study plus 29 globally sequences (randomly sampled) from all lineages 1A, 1B, 2, 3, 4, 5, 7, 8 (**Table S2**)**.** Sequences were aligned using MAFFT [13] and edited using AliView [14]. Those datasets were assessed for presence of phylogenetic signal by applying the likelihood mapping analysis implemented in the IQ-TREE 1.6.8 software [15]. A maximum likelihood phylogeny was reconstructed using IQ-TREE 1.6.8 software under the HKY+G4 substitution model [15]. We inferred time-scaled trees by using TreeTime [16].

**WNV epidemiological data**

Human reported cases presenting neurological disease, compatible with WNV infection collected between November 2015 and early 2020 were supplied by Brazilian Ministry of Health. We reinforce the nature of the reports as suspected (not confirmed), being officially defined as cases presenting neurological syndromes compatible with WNV infection. As such, the suspected cases time series should only be interpreted as a proxy for the possible geo-temporal dynamics of WNV infections. We restricted our analyses to Brazilian states for which we had sequence data - Minas Gerais, São Paulo and Piauí [17].

**Modelling transmission potential**

To estimate the transmission potential of WNV we employed the computational approach from Lourenço et al. recently applied in Israel [18]. This approach estimates the suitability index P using climatic variables only. The index measures the transmission potential of single adult female mosquitoes (spp. *Culex*) in the animal reservoir – and is thus interpreted as a summary measure of the risk for spill-over into human populations. The theory and practice of estimating the index P for mosquito-borne viruses has been previously described in full by Obolski et al. [19]. The epidemiological priors used were the same as in the original study by Lourenço et al. in Israel (**see Table in main text**), which relate to spp. *Culex*, WNV and an average bird species.

Climatic data was obtained from Copernicus.eu (https://www.copernicus.eu), in particular we used the dataset “essential climate variables for assessment of climate variability from 1979 to present” [20]. This dataset offers climatic variables at a time resolution of 1 month and gridded spatial resolution of 0.25°x0.25°. The climatic data was used in two ways to estimate the index P: (1) we averaged the climatic variables across the spatial dimension of a state to obtain a time series of estimated transmission potential, reporting the mean and variation per month; (2) and we used the climatic variables per spatial cell to obtain a local time series of estimated transmission potential (similar to that presented in [19] for South America).

**Supplementary Figures**

**Supplementary Figure 1.** WNV typing tool.

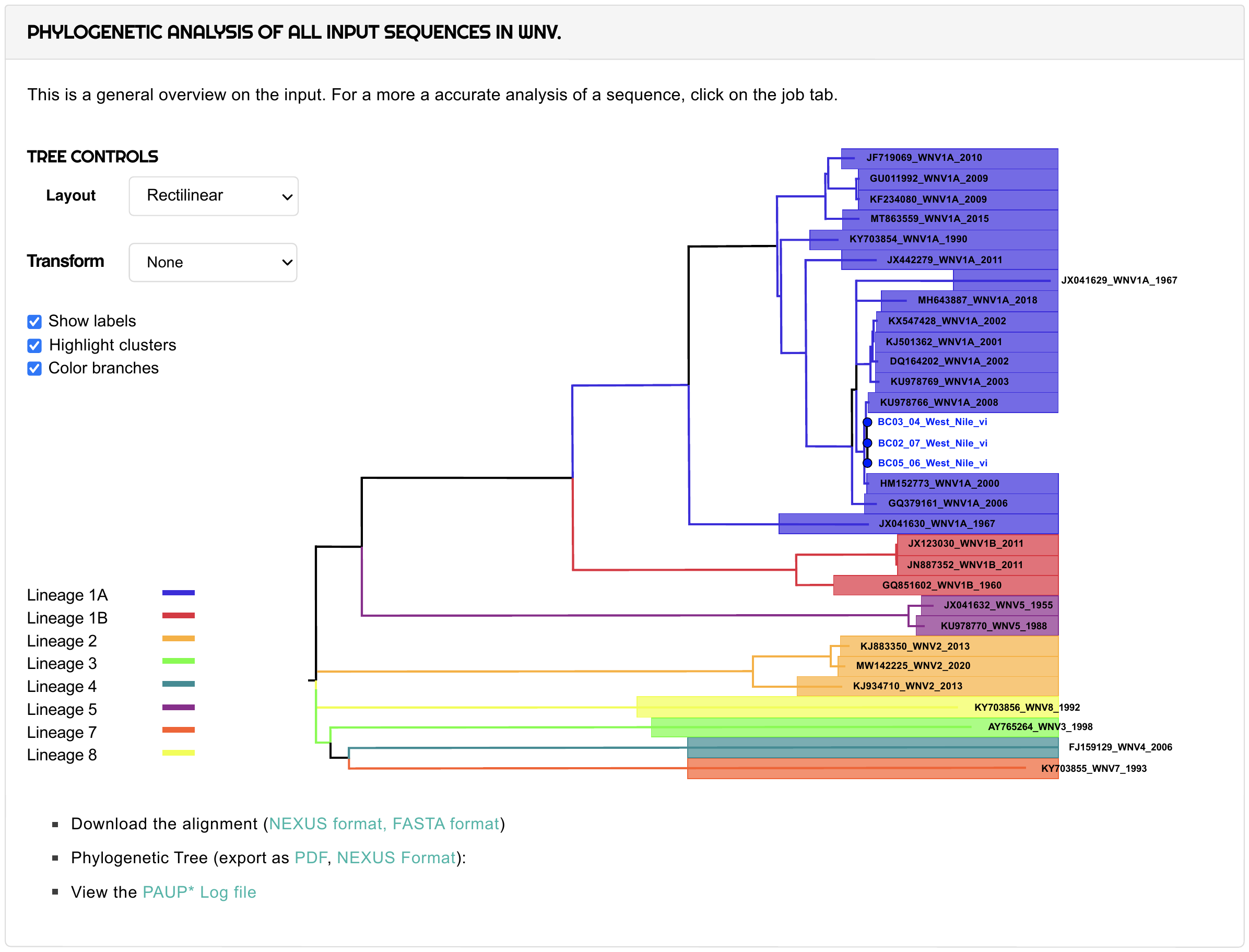

**Supplementary Figure 2.** Maximum likelihood phylogenetic tree of 2321 WNV complete genomes. Colours indicates different lineages. Highlighted red clade include the WNV viral strain obtained in this study.

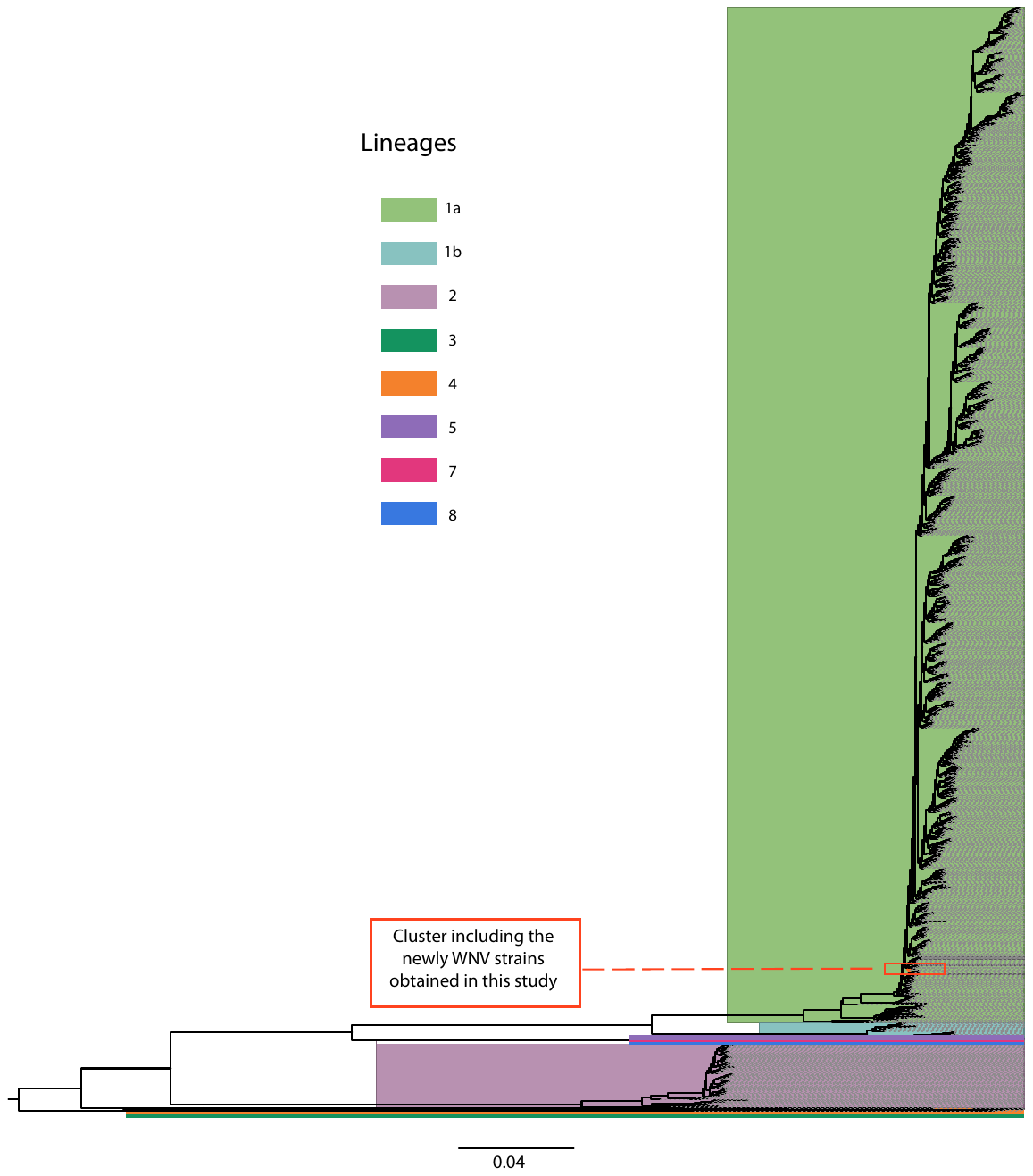

**Supplementary tables**

**Supplementary Table 1**

**Primer scheme**

| Primer Name | Sequence |
| --- | --- |
| WNVL1a_1_LEFT | GCCTGTGTGAGCTGACAAACTT |
| WNVL1a_1_RIGHT | TTTGTTTTGAGCTCCGCCGATT |
| WNVL1a_2_LEFT | GGATCGGTGGAGAGGTGTGAAT |
| WNVL1a_2_RIGHT | GACTTTGTGCACCAACAGTCGA |
| WNVL1a_3_LEFT | TGTCAGAGCAATGGATGTGGGA |
| WNVL1a_3_RIGHT | CTGTTGCTCATTCCAAGGCAGT |
| WNVL1a_4_LEFT | GTCATTGGTTGGATGCTTGGGA |
| WNVL1a_4_RIGHT | TGTGTCAATGCTTCCTTTGCCA |
| WNVL1a_5_LEFT | AATGACAAACGTGCTGACCCAG |
| WNVL1a_5_RIGHT | CACTCACGATGGACCAAGAACG |
| WNVL1a_6_LEFT | GGAGAATATGGAGAAGTGACAGTGG |
| WNVL1a_6_RIGHT | AAAGCCTTTGAACAGACGCCAT |
| WNVL1a_7_LEFT | TTCAAGCAACACTGTCAAGTTGAC |
| WNVL1a_7_RIGHT | TTTCCCAATGCTGCTTCCAGAC |
| WNVL1a_8_LEFT | CCTGATTGAATTGGAACCACCCT |
| WNVL1a_8_RIGHT | ACGGAGAGGAAGAGCAGAACTC |
| WNVL1a_9_LEFT | GCATGTCCTGGATAACGCAAGG |
| WNVL1a_9_RIGHT | AACCACGACACTAAGGTCCACA |
| WNVL1a_10_LEFT | TCTACGATCAGTTTCCAGACTGGA |
| WNVL1a_10_RIGHT | GTTGTTCTTGACAGCCGTTCCA |
| WNVL1a_11_LEFT | AGTGGAGGATTTTGGATTTGGTCT |
| WNVL1a_11_RIGHT | AGTCAATCTCTACCCGGCCTTC |
| WNVL1a_12_LEFT | GGCGATGGAATCCTTGAGAGTG |
| WNVL1a_12_RIGHT | GGCCCAACTGAAAAGGGTCAAT |
| WNVL1a_13_LEFT | CGGCTGTTGGTATGGTATGGAG |
| WNVL1a_13_RIGHT | GAAAACAGCCGCCAACATCAAC |
| WNVL1a_14_LEFT | GGCGACCTTCAAGATACAACCA |
| WNVL1a_14_RIGHT | GCTAGAGCCAAGCATAGCAGAC |
| WNVL1a_15_LEFT | GGATACTGCTGTTGATGGTCGG |
| WNVL1a_15_RIGHT | TCATCAAGCCGCACATCAACTC |
| WNVL1a_16_LEFT | TTCTGGGAAATCAACAGATATGTGGA |
| WNVL1a_16_RIGHT | TCAAAGCGGCTCCTTTTGTTGT |
| WNVL1a_17_LEFT | TCTACAGGATCATGACTCGCGG |
| WNVL1a_17_RIGHT | CGCTTATGTATGAGCCGTTGGG |
| WNVL1a_18_LEFT | GACTTTGGACTTCCCCACTGGA |
| WNVL1a_18_RIGHT | ATTATGTTCTCTGGGCACTGCG |
| WNVL1a_19_LEFT | ACAGAAGACTGAGAACAGCCGT |
| WNVL1a_19_RIGHT | TCCAGAGTTCCAAGCTCGATCC |
| WNVL1a_20_LEFT | TATTCATGACAGCCACCCCACC |
| WNVL1a_20_RIGHT | GCTGTCACTGCAGATGGTTCTC |
| WNVL1a_21_LEFT | CTAACTTCAAGGCGAGCAGGGT |
| WNVL1a_21_RIGHT | TGCAGTCCTCAACAGTTCCAGA |
| WNVL1a_22_LEFT | TCTACCAACCAGAGCGTGAGAA |
| WNVL1a_22_RIGHT | CCCAGAACCTCAATGAGCCCTA |
| WNVL1a_23_LEFT | GAAAGGAAGATTCTGAGGCCGC |
| WNVL1a_23_RIGHT | CGTTCCTGGAACTTCAGCCATC |
| WNVL1a_24_LEFT | GTATTCTTCCTCCTCATGCAGCG |
| WNVL1a_24_RIGHT | CGGCCTCAAGTCCAGAAGAAAC |
| WNVL1a_25_LEFT | GTTGGCTGGACAAGACCAAGAG |
| WNVL1a_25_RIGHT | GGAACCATGTAGGCATAGTGGC |
| WNVL1a_26_LEFT | CTTCGTCGATGTTGGAGTGTCG |
| WNVL1a_26_RIGHT | CTCCATTCTCCCAAAGCGTCAC |
| WNVL1a_27_LEFT | GCTGATCTTAGTGTCTCTAGCTGC |
| WNVL1a_27_RIGHT | CAGTTTTGCTGTGCCCCTAGAG |
| WNVL1a_28_LEFT | GTACCGCAAAGAGGCCATCATC |
| WNVL1a_28_RIGHT | TTGACGAGGACTCTCCGATGTC |
| WNVL1a_29_LEFT | TGGAACATTGTCACCATGAAGAGT |
| WNVL1a_29_RIGHT | CTTCCTCGTATTGGGGTCCCTT |
| WNVL1a_30_LEFT | GAGTCGAGCTTCAGGCAATGTG |
| WNVL1a_30_RIGHT | AGGGAGTAGTGTCAGTCATGGC |
| WNVL1a_31_LEFT | ACGGCAGTTATGATGTGAAGCC |
| WNVL1a_31_RIGHT | CTCCTCATCCACCATCTCCCAA |
| WNVL1a_32_LEFT | GTCAACAGCAATGCAGCTTTGG |
| WNVL1a_32_RIGHT | GCCAACTTCACGCAGGATGTAA |
| WNVL1a_33_LEFT | TCGAGGCTCTGGGTTTTCTCAA |
| WNVL1a_33_RIGHT | TTCCCCTTCCATCATCCTCACC |
| WNVL1a_34_LEFT | TCCAGAGAAGATCAGAGGGGGA |
| WNVL1a_34_RIGHT | TGGAACCACCAGTGTTCTTCCA |
| WNVL1a_35_LEFT | AGAGTGGAAACCGTCAACTGGA |
| WNVL1a_35_RIGHT | CCAGACCTCCAACATGTCCTCT |
| WNVL1a_36_LEFT | GTGGCTGCTTCTGTACTTCCAC |
| WNVL1a_36_RIGHT | TCTACAGTACTGTGTCCTCAACCA |
| WNVL1a_37_LEFT | GTGGCTATCAACCAAGTCAGAGC |
| WNVL1a_37_RIGHT | CAACATGTGGGGTCCTTCTTCC |
| WNVL1a_38_LEFT | GAAGTTGAGTAGACGGTGCTGC |
| WNVL1a_38_RIGHT | ACGGGGTCTCCACTAACCTCTA |

**Supplementary Table 2.** Globally reference WNV sequences from the subset n=29 used in this study

| **Acession Number** | **Collection date** | **Country** | **Lineage** | **Host** |
| --- | --- | --- | --- | --- |
| MN849176 | 01/09/2019 | USA | 1a | Mosquito |
| JF719069 | 01/08/2010 | Spain | 1a | Horse |
| KY703854 | 01/01/1990 | Senegal | 1a | Mosquito |
| GU011992 | 01/01/2009 | Italy | 1a | Human |
| KF234080 | 01/01/2009 | Italy | 1a | Human |
| MT863559 | 10/03/2015 | France | 1a | Horse |
| JX442279 | 01/01/2011 | China | 1a | Mosquito |
| JX041629 | 01/01/1967 | Azerbaijan | 1a | Bird |
| JX041630 | 01/01/1967 | Azerbaijan | 1a | Bird |
| KX547428 | 26/09/2002 | USA | 1a | Mosquito |
| KJ501362 | 01/01/2001 | USA | 1a | Crow |
| DQ164202 | 01/01/2002 | USA | 1a | Human |
| MH643887 | 26/04/2018 | Brazil | 1a | Horse |
| KU978769 | 30/05/2003 | Mexico | 1a | Crow |
| KU978766 | 01/07/2009 | Colombia | 1a | Flamingo |
| GQ379161 | 01/02/2006 | Argentina | 1a | Horse |
| HM152773 | 01/01/2000 | Israel | 1a | Human |
| JX123030 | 03/07/2011 | Australia | 1b | Horse |
| GQ851602 | 01/01/1960 | Australia | 1b | Mosquito |
| JN887352 | 01/01/2011 | Australia | 1b | Horse |
| KJ883350 | 05/07/1905 | Greece | 2 | Human |
| KJ934710 | 27/08/2013 | Romania | 2 | Tick |
| MW142225 | 01/08/2020 | Germany | 2 | Human |
| AY765264 | 01/01/1998 | Czech Republic | 3 | Mosquito |
| FJ159129 | 01/01/2006 | Russia | 4 | Mosquito |
| JX041632 | 01/01/1955 | india | 5 | Mosquito |
| KU978770 | 02/12/1988 | india | 5 | Human |
| KY703855 | 01/01/1993 | Senegal | 7 | Tick |
| KY703856 | 01/01/1992 | Senegal | 8 | Mosquito |

**References**

[1] Stanislawek WL, Blacksell SD, Newberry KM, et al. Detection by ELISA of bluetongue antigen directly in the blood of experimentally infected sheep. Vet Microbiol. 1996;52:111-112.

[2] Fulop L, Barrett ADT, Phillpotts R, et al. Rapid identification of flaviviruses based on conserved NS5 gene sequences. J Virol Methods. 1993; 44:179–188.

[3] Pfeffer M, Proebster B, Kinney RM, et al. Genus‐specific detection of alphaviruses by a semi‐nested reverse transcription‐polymerase chain reaction. Am J Trop Med Hyg. 1997;57:709–718.

[4] Silva ASG, Matos ACD, Cunha MACR, et al. Transbound Emerg Dis. 2019; 66:445-453.

[5] Costa EA, Rosa R, Oliveira TS, et al. Molecular characterization of neuropathogenic Equine Herpesvirus 1 Brazilian isolates. Arq Bras Med Vet. Zootec. 2015;67:1183–1187.

[6] Sorg I, Metzler A. Detection of Borna Disease Virus RNA in Formalin-Fixed, Paraffin-Embedded Brain Tissues by Nested PCR. J Clin Microbiol. 1995;4:821-823.

[7] Faria NR, Quick J, Claro IM, et al. Establishment and cryptic transmission of Zika virus in Brazil and the Americas. Nature. 2017;546:406-10.

[8] Quick J, Grubaugh ND, Pullan ST, et al. Multiplex PCR method for MinION and Illumina sequencing of Zika and other virus genomes directly from clinical samples. Nat Protoc. 2017;12:1261-76.

[9] Vilsker M, Moosa Y, Nooij S, et al. Genome Detective: an automated system for virus identification from high-throughput sequencing data. Bioinformatics. 2019; 35: 871–3.

[10] Altschul SF, Gish W, Miller W, et al. Basic local alignment search tool. Journal of Molecular Biology. 1990;215:403-10.

[11] Larkin MA, Blackshields G, Brown NP, et al. Clustal W and Clustal X version 2.0. Bioinformatics. 2007;23:2947-8.

[12] Lemey P, Salemi, M., Vandamme, A. The Phylogenetic Handbook: A Practical Approach to Phylogenetic Analysis and Hypothesis Testing. 2 ed. Cambridge: Cambridge University Press; 2009.

[13] Katoh K, Kuma K, Toh H, et al. MAFFT version 5: improvement in accuracy of multiple sequence alignment. Nucleic Acids Res. 2005; 33:511–8.

[14] Larsson A. AliView: a fast and lightweight alignment viewer and editor for large data sets. Bioinformatics. 2014;30:22-2.

[15] Nguyen LT, Schmidt HA, von Haeseler A, Minh BQ. IQ-TREE: a fast and effective stochastic algorithm for estimating maximum-likelihood phylogenies. Mol Biol Evol. 2015;32:268-74.

[16] Sagulenko P, Puller V, Neher RA. TreeTime:Maximum-likelihood phylodynamic analysis. Virus Evol. 2018;4:42-44.

[17] Ministério da Saúde (MS). Secretaria de Vigilância em Saúde. Monitoramento da Febre do Nilo Ocidental no Brasil, 2014 a 2019 (Nota Informativa). Brasília: MS; 2019. 7p.[https://antigo.saude.gov.br/images/pdf/2019/julho/08/informe-febre-niloocidental-n1-8jul19b.pdf]

[18] Lourenço J, Thompson RN, Thézé J, et al. Characterising West Nile virus epidemiology in Israel using a transmission suitability index. Euro Surveill. 2020;2:5-41.

[19] Obolski U, Perez PN, Villabona-Arenas CJ, et al. MVSE: An R-package that estimates a climate-driven mosquito-borne viral suitability index. Methods Ecol Evol. 2019;10:1357–1370.

[20] Copernicus Climate Data Store. [cited 23 Dec 2020]. Available: <https://cds.climate.copernicus.eu/cdsapp#!/dataset/ecv-for-climate-change?tab=overview>.
